# Primate ventromedial prefrontal cortex neurons continuously encode the willingness to engage in reward directed behavior

**DOI:** 10.1101/062315

**Authors:** Aurore San Galli, Chiara Varazzani, Raphaelle Abitbol, Mathias Pessiglione, Sebastien Bouret

## Abstract

To survive in their complex environment, primates must integrate information over time and adjust their actions beyond immediate events. The underlying neurobiological processes, however, remain unclear. Here, we assessed the contribution of the ventromedial prefrontal cortex (VMPFC), a brain region important for value-based decision making. We recorded single VMPFC neurons in monkeys performing a task where obtaining fluid rewards required squeezing a grip. The willingness to perform the action was modulated not only by visual information about Effort and Reward levels, but also by contextual factors such as Trial Number (i.e fatigue and/or satiety) or behavior in recent trials. A greater fraction of VMPFC neurons encoded contextual information, compared to visual stimuli. Moreover, the dynamics of VMPFC firing was more closely related to slow changes in motivational states driven by these contextual factors rather than rapid responses to individual task events. Thus, the firing of VMPFC neurons continuously integrated contextual information and reliably predicted the monkey’s willingness to perform the task. This function might be critical when animals forage in a complex environment and need to integrate information over time. Its relation with motivational states also resonates with the VMPFC implication in the default mode or in mood disorders.

## Introduction

Natural selection puts a strong pressure on all animals to optimize the ratio between reward-associated costs and benefits (Altman, 2006; Milton and May, 1976; Stephens and Krebs, 1986). This optimization can be achieved by simple stereotyped reflexes to behaviorally relevant stimuli (Berridge, 2004; Lorenz, 1981). With only these reflexes, however, behavior would remains directly bound to the immediate environment, and in several species goal-directed behavior enables to integrate information over space and time (Aminoff et al., 2013; Balleine and Dickinson, 1998; Clayton et al., 2003; Correia et al., 2007; Fuster, 2008). In primates, this would be crucial since most of them are frugivorous and fruiting trees are both sparsely distributed and highly seasonal: they could not just wander randomly and expect to find fruits ‘by chance’ in the forest (Janmaat 2011; 2013; Cunningham, 2007; Noser, 2015; Cunningham and Janson 2007). This ecological pressure might have driven the evolution of specific cognitive abilities associated with the prefrontal cortex, which is particularly developed in primates (Fuster, 2008; Genovesio et al., 2013; Passingham et al., 2012).

Recent work has emphasized the key role of the ventral prefrontal cortex in the representation of reward values (Boorman et al., 2013; Bouret and Richmond, 2010; Chib et al., 2009; Hosokawa et al., 2013; Kable and Glimcher, 2007; Klein-Flugge et al., 2013; Lebreton et al., 2009; O'Doherty et al., 2001; Padoa-Schioppa and Assad, 2006; Strait et al., 2014; Walton et al., 2009). Anatomically, the medial and orbital regions can be dissociated by the strength of connections with the hippocampus and parahippocampal cortices (stronger for the medial network) and sensory structures (stronger for the orbital network) (Lavenex and Amaral, 2000; Ongür and Price, 2000). Functionally, however, this distinction remains debated. In humans, there is a consensus regarding the specific implication of the ventromedial prefrontal cortex (VMPFC) in processing subjective value (Bartra et al., 2013; Clithero and Rangel, 2014; Kable and Glimcher, 2007; Lebreton et al., 2009; Rangel et al., 2008; Rushworth et al., 2011). In monkeys, activity related to reward value has been traditionally described in the orbitofrontal cortex, especially when reward information is provided by sensory stimuli (Padoa-Schioppa and Assad, 2006; Roesch and Olson, 2004; Thorpe et al., 1983; Tremblay and Schultz, 2000). However, more recent studies indicate that the VMPFC may also encode value in monkeys, especially when it relies upon internal information (Abitbol et al., 2015; Bouret and Richmond, 2010; Noonan et al., 2010; Strait et al., 2014). This is coherent with studies emphasizing the importance of the interaction between VMPFC and the medial temporal lobe to attribute value to imaginary items (Barron et al., 2013; Benoit et al., 2014; Clark et al., 2013; Lebreton et al., 2013; Peters and Büchel, 2010).

In that framework, the primate VMPFC should be critical for adjusting the willingness to engage in the current course of action based on a slow accumulation of internal and external information. Critically, this estimate should not be limited to specific events implemented in experimental tasks. Rather, it should continuously integrate information over time, irrespectively of its source (sensory stimuli or contextual information). To test this hypothesis, we recorded single unit activity in monkeys performing a task where reward and effort levels were systematically manipulated. We focused on area 14r, a part of the VMPFC which is specific to primates (Wise, 2008). In line with our hypothesis, we found that VMPFC activity was related to several factors driving the willingness to engage in the task. Importantly, this relation was both slow and coherent over time, in line with the idea that VMPFC activity reflects states of motivation rather than discrete event-related functions. These results are in line with recent studies in humans indicating that the role of the VMPFC in value-based decision making is especially critical for guiding behavior based on evaluations processes reaching beyond the immediate environment.

## Material & Methods

### Animals

We used 2 sub-adults male macaque monkeys for these experiments, A (4 years, 6 kg) and B (5 years, 7 kg kg). Monkeys were housed in a group of 6 individuals, with free access to food and controlled access to water during the course of the experiments. Monkeys received water as a reward for performing the task. Experiments were carried out in accordance with the European Community Council Directive and the french legislation (Ministère de l’Agricul^−^ ture et de la Forêt, Commission nationale de l’expérimentation animale) (86/609/EEC)). They were approved by the Darwin ethics comity of the university Paris 6 (CREEA IDF n° 3).

### Behavior

The behavioral setting was identical to that of our recent study (Varazzani et al., 2015). Each monkey squatted in a primate chair positioned in front of a monitor on which visual stimuli were displayed. A pneumatic grip (M2E Unimecanique, Paris, France) was mounted on the chair at the level of the monkey’s hands. Liquid rewards were delivered from a tube positioned between the monkey’s lips. Eye position and pupil area were monitored continuously using a video-based eye tracker (Iscan inc, MA, USA). The behavioral paradigm was controlled using the REX system (NIH, MD, USA) and Presentation software (Neurobehavioral systems, Inc, CA, USA)).

Monkeys were trained to perform a simple force task: A red target point (wait signal) appeared at the center of the monitor. After a random interval of 500-1500 ms, the target turned green (go signal). If the monkey squeezed the grip 200-1000 ms after the green target appeared, the target turned blue (feedback) and the monkey had to maintained the effort for another 300-600 ms in order to get the fluid reward. For this initial phase, the required effort was adjusted to approximately 70% of the maximum force, assessed by progressively increasing the threshold necessary to obtain the reward.

Once monkeys were comfortable with this task, we progressively introduced the 2 parameters of interest (3 levels of effort and 3 levels of reward) as well as the corresponding visual cues. Importantly, error trials were repeated, to prevent monkeys from systematically ‘skipping’ the least favorite conditions. The cue appeared within 1 second after the onset of the red ‘wait’ signal and remained on the screen until the end of the trial. We used several cue sets per monkey, mostly during the initial recording but also during the recording sessions. All cues were grey-levels isoluminant fractals or scrambled versions of the same original fractal image, in order to minimize luminance differences (Figure 1A). Monkeys were trained until they could reliably express a differential behavior across the 9 conditions.

**Figure 1:**
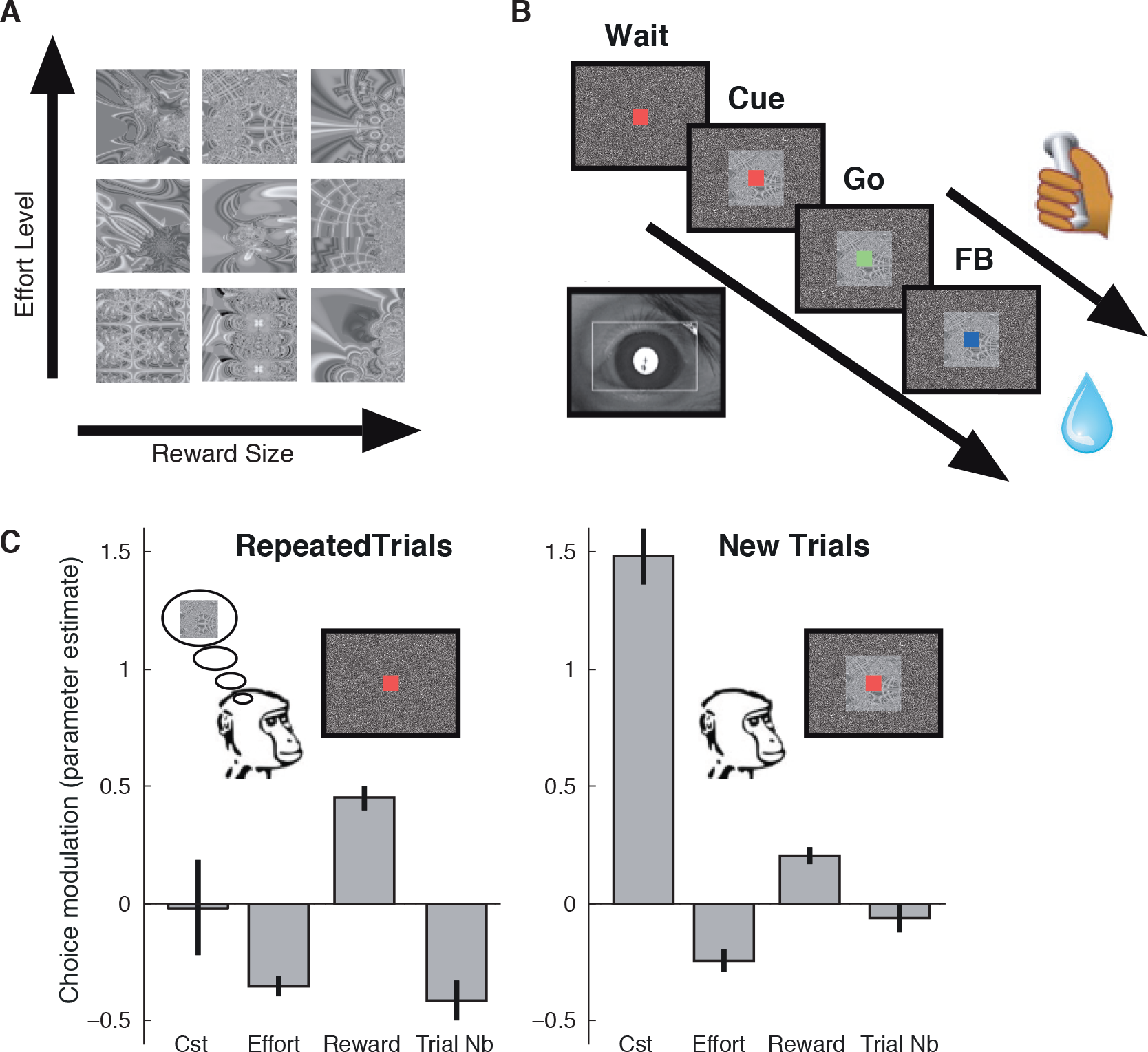
*Task and behavior*. *Monkeys performed an operant task where they must exert a physical force (squeezing a grip) to obtain fluid reward. **A**) There were 9 trial types, defined by a combination of 2 factors, each with 3 levels: Reward Size (1, 2 or 4 drops) and Effort Level (low, medium or high). Each trial was signaled to the monkey using a specific visual cue. **B**) Trials started with the onset of a red fixation point (‘**wait**’). Monkeys must keep their gaze on that central spot during the entire duration of the trial, or it was aborted. Within 500-800 ms after the onset of the fixation point, a visual cue appeared to indicate which trial type the monkey was in (‘**Cue**’). Within 1-2 seconds after cue onset, the fixation point turned green (go signal), indicating that the monkey must squeeze the bar with the cued amount of force, within 1 second (‘**Go**’). The fixation point turned blue (feedback) when they reached the required force (**FB**). Monkeys could overshoot if they wanted too, all they had to do was to maintain the exerted force above the required level until the reward was delivered (random delay of 200-400 ms). If monkeys made an error at any step of the trial, an error was scored and the trial was repeated until it was performed correctly. **C**) Influence of Effort Level, Reward Size, and Trial Number on the choice to perform the trial in New trials (right), where the information is provided by visual cues, and in Repeated trials (left), where information is provided by memory (not necessarily of the cue itself). We assessed the influence of the 3 factors, plus a constant, on choices using a logistic regression to estimate their respective coefficient. Parameter estimates are represented as the mean ± SEM of the regression coefficients across all sessions. Data from the 2 monkeys did not differ so they were pooled for clarity. In both situations, the choice to perform the trial was influenced positively by the size of the expected reward and negatively by the effort level and progression through the session. Monkeys made globally more positive choices in New trials (higher constant), and choices were globally less sensitive to task factors (smaller coefficients)*.

Once monkeys had reached a stable performance, a 3 T MR image was obtained to determine the location of the ventromedial prefrontal cortex to guide recording well placement. Then, a sterile surgical procedure was carried out under general isoflurane anesthesia in a fully equipped and staffed surgical suite to place the recording well and the head fixation post. Training resumed 4 weeks after surgery.

Monkeys were trained to perform the task with the head fixed, and then trained to fixate a central spot. Following these simple procedures, we introduced the final version of the task, which required the monkey to fixate the ‘wait’ (red) spot for 500-800 ms before the cue appeared. Monkeys were required to fixate the central spot until reward delivery. The go signal (i.e. red point turned green) appeared 800-1600 ms after cue onset, and the monkey had up to 1 sec to respond by squeezing the grip. Once they had reached the required level of force, the point turned blue and the monkey had to maintain the effort level above the required threshold for another 300-600 ms in order to get the reward. The inter-trial interval lasted 1300-1700 ms. Again, error trials were repeated.

We sorted the types of errors in 2 categories, those reflecting a choice of the animal to not perform the trial (fixation break and omissions to squeeze the bar) and those reflecting a failed attempt to perform the task (when the exerted force did not reach the required threshold or when the monkey released the grip too early). The latter (around 1% of the trials) were considered together with correct trials as choices to perform the trial. In other words, a choice was counted as positive when monkeys squeezed the bar, and all other cases were counted as negative choices. In the vast majority of cases, monkeys decided to forgo the trial by breaking fixation, which could happen before or after cue onset.

### Electrophysiology

Electrophysiological recordings were made with tungsten microelectrodes (FHC, impedance: 1.5 MΩ). The electrode was positioned using a stereotaxic plastic insert with holes 1mm apart in a rectangular grid (Crist Instruments). The electrode was inserted through a guide tube. After several recording sessions, MR scans were obtained with the electrode at one of the recording sites; the position of the recording sites was reconstructed based on relative position in the stereotaxic plastic insert and on the alternation of white and grey matter based on electrophysiological criteria during recording sessions. All recording sites were located in the ventral part of the girus rectus, from 1 to 7 mm anterior to the genu of the corpus callosum. All recordings were obtained from area 14r, based on architechtonic maps of Carmichael & Price (1994). All the neurophysiological and behavioral data were collected on an Omniplex system (Plexon Co, Tx, USA). The signal was amplified (× 10 000) and filtered (100 Hz-2 kHz) for single unit sorting. Single units were isolated offline using plexon software.

### Data analysis

To estimate the willingness to work on a trial by trial basis, we measured the choice to perform the action, or not, as a function of task parameters (Reward and Effort levels) and contextual effects including progression through the session and past responses. The choice variable was equal to 1 in trials where monkeys squeezed the bar, and otherwise it was equal to zero. Note that in some trials monkeys did not even fixate the point long enough to let the cue appear, but modulated their behavior based on contextual information. We used a logistic regression to predict trial by trial choices based on a constant term and several parameters (e.g. Effort and Reward levels). We estimated the regression coefficients of each of these parameters using the *glmfit.m* function in Matlab, with a logit link. We used this measure to evaluate the willingness to work either in specific subsets of trials (Repeated trials and New trials) or in all trials, whether or not monkeys fixated long enough to let the cue appear. In the later case, the variable ‘information about Reward’ or ‘information about Effort’ was scored as 0 when it was not available (e.g. in New trials, if monkeys broke fixation before the cue appeared). In all other cases (in Repeated trials or in New trials if monkeys fixated long enough to let the cue appear before choosing to respond or not), the monkeys readily had access to the information about Reward and Effort. Repeated trials correspond to trials where monkeys failed to complete the previous trials but thereby obtained information about Force and Reward levels for the current one. Since monkeys are familiar with the structure of the task (erroneous trials are repeated), they can infer that Reward and Effort will be the same as in the previous trial, choose to engage in the trial or not from the onset of the wait signal, before the cue appears. New trials were trials following a correct response, where Reward and Effort levels could only be inferred from the visual cues. To estimate the progression through the session and the resulting effect of fatigue and satiety, we measured the cumulated sum of trials since the beginning of the recording session (Bouret & Richmond, 2010). We also examined the influence of responses in past trials in a systematic fashion, over distances ranging from 1 to 35 trials back. Importantly, we systematically included potential confounding factors (Reward, Effort and Trial Number) as co-regressors in the model, to evaluate the effect of past responses on current behavior over and above these parameters.

We used a similar approach to evaluate the influence these factors on neuronal activity. We measured the firing rate of each VMPFC neurons in 3 windows: (1) from 0 to 500 ms after the onset of the fixation point, but only in repeated trials; (2) from 100 to 600 ms after cue onset, but only in unrepeated trials; (3) from 0 to 500 ms after the onset of reward delivery. To measure the neural encoding of task factors, we used Generalized Linear Model (GLM) in which neuronal single-trial firing rates were modeled as a constant factor plus a weighted linear combination of three variables: Effort level, Reward size and the Trial number. We used the estimated regression coefficients of each of these variables to compare their relative influence on neuronal activity. The variables were z-scored for each neuron to allow comparison of the effects. The firing rates were raw data expressed in spikes per second. We also used a GLM to estimate the relation between firing and willingness to work, over and above other factors such as task parameters (by including them as co-regressors)

To compare the relation between VMPFC firing and engagement in the task across several time scales, we measured the firing rate and the number of trials completed in bins of several sizes. For each bin size, the entire session was split into successive bins in which we counted not only the firing rate (number of spikes fired by the neuron in each bin) and the work rate (number of trials completed in each bin), but also the average reward size and the average effort size over all the trials within the bin. Indeed, slow fluctuation in VMPFC activity and Work Rate could be directly driven by slow changes in Reward/ Effort or by the progression through the session, so we needed to estimate the variance explained by slow fluctuations above and beyond these factors. Practically speaking, we ran 2 analyses: in the first one, we simply computed the correlation between work rate and firing rate across all bins. In the second model of firing rate, task variables (Reward, Effort and Trial number) were included as coregressors, along with Work Rate, from which we regressed out the variance due to task factors (reward, effort and trial number). Thus, for each bin size, we tried to predict VMPFC firing rates using a GLM to estimate the coefficients of 4 parameters: Work Rate (corrected), average Reward, average Effort, Bin number, plus a constant term.

## Results

### Behavior

We trained two monkeys to perform the effort/reward task depicted in Figure 1A-B. First, monkeys readily used the information about upcoming effort to adjust their behavior. We assessed the influence of Effort level on the amount of force produced using a linear regression, which was significant in both animals: monkey A: β_E_;=0.8 ±0.009, p<10^−3^; monkey B: β_E_;=0.9 ±0.004, p<10^−3^. Thus, monkeys produced the minimum amount of effort required to complete the trial, rather than producing a fixed high force.

Second, monkeys did not systematically perform the action to obtain the reward: on average, monkey A accepted to perform the task in 52 ± 5% of the trials (22 sessions) and monkey B accepted in 66 ± 4% of the trials (36 sessions). In this task, since trials were repeated until they were performed correctly, monkeys could choose to engage in the trial in two very distinct conditions: Repeated and New trials. In New trials, which followed a correctly performed action, information about upcoming Reward and Effort was provided by the visual cue. By contrast, in Repeated trials, upcoming Reward and Effort levels could be inferred using general knowledge about the task structure (error trials are repeated) and memory of the previous trial (either the cue itself, the information, or the decision). Thus at the trial onset (fixation point), monkeys could choose to engage in the trial, or not, based on information in memory rather than based on visual cues. Hence, we separated choices made at the onset of the fixation point in Repeated trials and choices made after the cue onset in New trials. In both cases, the choices consisted either in maintaining fixation and perform the action (‘yes’), or in breaking fixation and abort the trial (‘no’) (see methods for further details).

Both monkeys modulated their choices to perform the action as a function of task parameters (the expected amount of reward and effort) and as a function of progression through the session (number of trials performed), which combines fatigue and satiety. But these effects differed between Repeated and New trials (Figure 1C). For each condition (Repeated vs New trials) and each monkey (A and B), we used a logistic regression to evaluate the influence of a constant plus 3 parameters (Reward, Effort, and Trial Number) on the choice to perform the action, or not. The interactions among the 3 task factors were not included because initial analysis showed that they were not significant. All regressors were z-scored to allow a direct comparison of the estimated coefficients. We then compared these coefficients using a 3 way ANOVA, to evaluate the influence of 3 factors: Monkey (2 categories, one for each animal); Condition (2 categories, Repeated vs New trials) and Variable (3 categories: Reward Size, Effort Level and Trial Number). There was no difference between the 2 animals (no main effect and no interaction involving the factor Monkey was significant (all p>0.1). As shown on Figure 1C, the constant was greater in New trials, indicating that globally monkeys were more willing to perform the action in those trials, compared to repeated trials. Indeed, monkeys responded more positively in New (78 +/− 2%; average across animals) compared to Repeated trials (50 +/− 3%; average across animals). This is presumably because at the time of the cue in New trials, monkeys had already committed to see the cue and obtain the information about costs and benefits. Thus, since they were already engaged, they probably had a bias for performing the action and they were less sensitive to information about costs and benefits. In line with this interpretation, the effects of all 3 factors (Effort, Reward and Trial Number) were greater in Repeated compared to New trials. First, there was a greater variability in behavior in Repeated trials (total variance= 0.24) compared to New trials (variance= 0.17). As expected, there was a clear difference among the effects of the 3 task variables (Main effect of Variable, F=65.6; p<10^−4^). Both Effort and Trial Number had a negative influence on choices, whereas Reward had a positive effect. But the greater variability in Repeated trials was captured by the clear interaction between the factors Variable and Condition (F=14, p<10^−4^). As shown on Figure 1C, the effects of all 3 factors were more pronounced in Repeated compared to New trials. In short, monkeys adjusted their willingness to engage in the task across conditions, defined by a combination of Reward and Effort level and the progression through the task (Trial number). In New trials, monkeys were globally more likely to perform the action and less sensitive to task factors, presumably because they were already engaged in the trial.

### Electrophysiology

We recorded 56 and 57 neurons from the VMPFC of monkey A and B, respectively (see Figure 2). All neurons encountered along the tracks were included in the analysis, as long as the units were well isolated using a time-voltage threshold discrimination criterion and as long as these criteria remained stable throughout the session. All neurons were recorded from the gyrus rectus, between 1 and 9 mm anterior to the genu of the corpus callosum (area 14r, Carmichael & Price, 1994). The activity profiles were similar in the two animals, so the neuronal data (113 units) were pooled. The average firing rate of those neurons was 3.6 ± 0.6 spk/sec.

**Figure 2:**
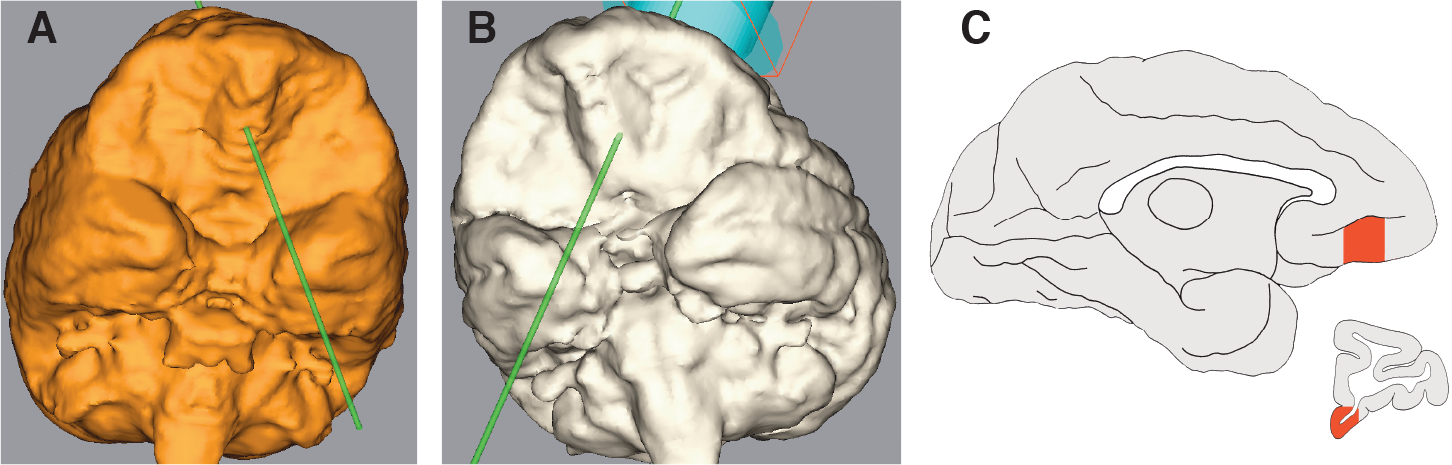
*Recording location.* *Panels **A** and **B** show a reconstruction of a typical electrode trajectory through the prefrontal cortex of monkey A and B, respectively. In both cases, the electrode was inserted in the middle part of the gyrus rectus. Panel **C** shows the area where the recordings were obtained in the 2 monkeys. This region corresponds to area 14r (Carmichael and Price, 1994).*

#### VMPFC neurons reliably encode reward size and trial number

We first examined the modulation of firing as a function of the 3 task variables, Reward Size, Effort Level and Trial Number. Figure 3 and Figure 4 show representative examples of single VMPFC neurons encoding Reward Size across task epochs. We used a GLM to estimate the influence of Effort, Reward and Trial Number on the firing rate of each neuron in sliding windows around 3 task events: i) cue onset in New trials (when information about reward and effort was provided by visual cues, Figure 5A), ii) onset of the fixation point in Repeated trials (when information about trial outcome could be inferred from the previous trial and knowledge about the task, Figure 5B), iii) reward delivery (the actual trial outcome, Figure 5C). All neurons were included in the analysis and all p-values were corrected for multiple comparisons across windows using an False Discovery Rate procedure.

**Figure 3:**
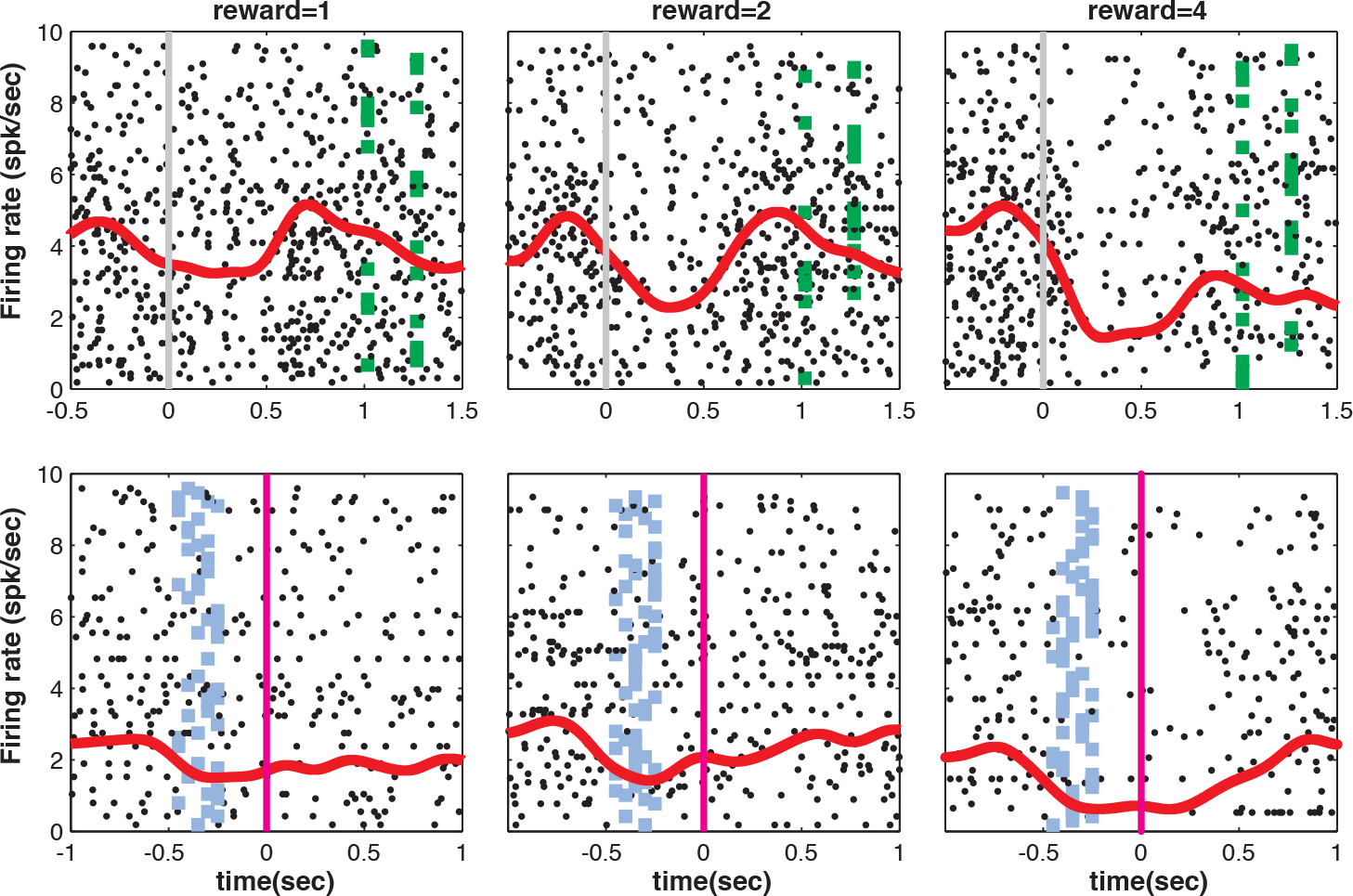
*Example of single neuron encoding reward size at two events of New trials*. *Spiking activity (raster display and cumulative spike density function, in red) of a representative VMPFC neuron around 2 task events: the onset of the cue (top, grey line) and the reward delivery (bottom, pink line). We also indicated the time of additional events in each trials: at the top, green dots represent the Go signal and at the bottom blue dots represent the onset of the feedback following correct actions. Both events were associated with a decrease in firing rate, which were more pronounced for high rewards (right) than for small rewards (left). Thus, the encoding of Reward by this neuron is very consistent over time*.

**Figure 4:**
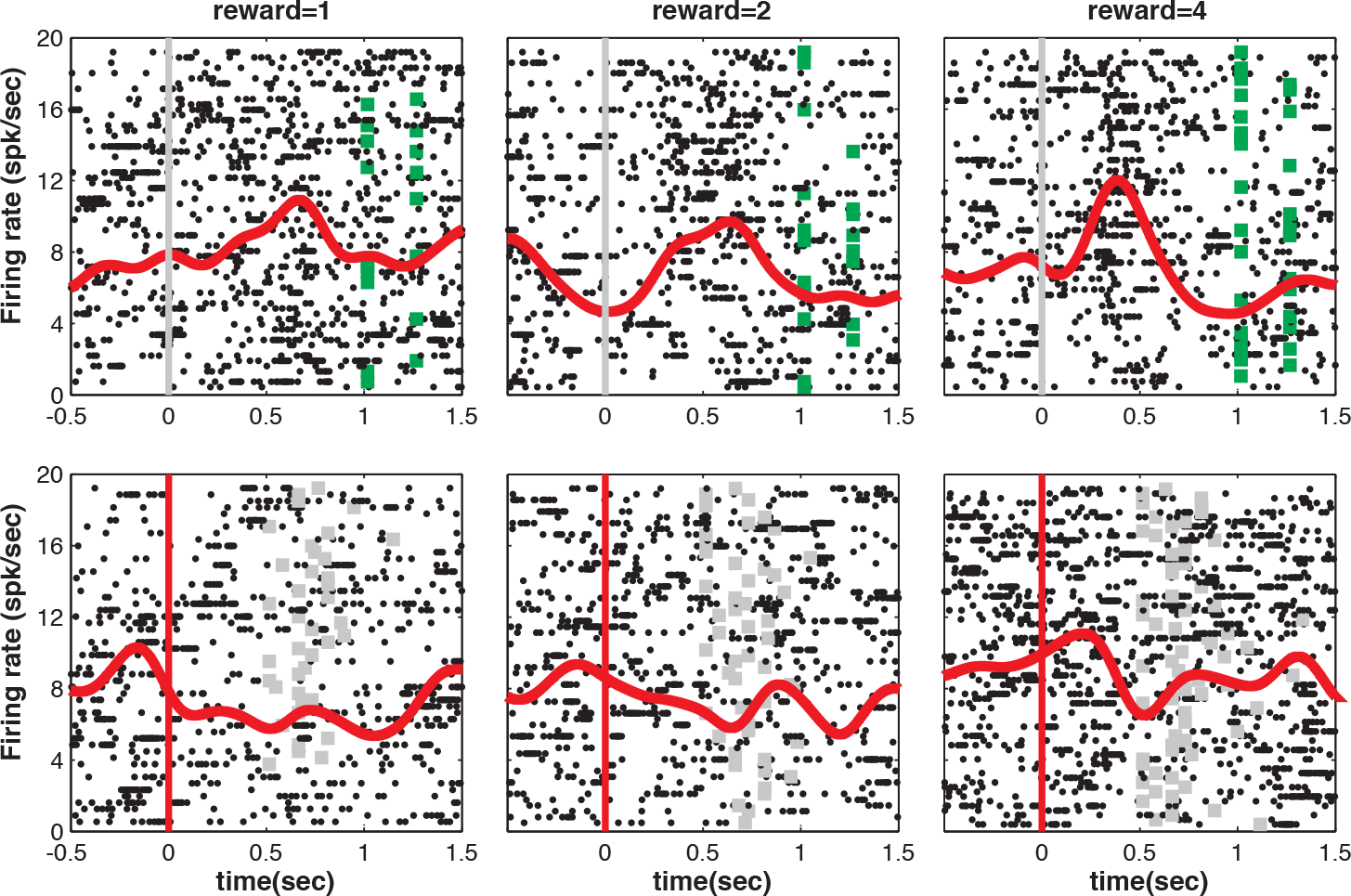
*Example of single neuron encoding reward size in New and Repeated trials*. *Spiking activity (raster display and cumulative spike density function, in red) of a representative VMPFC neuron around 2 task events: the onset of the cue in New trials, when reward information was provided by visual cues (top) and the onset of the fixation point in repeated trials, when reward information was in memory (bottom). At the top, green dots indicate the onset of the Go signal. At the bottom, grey dots indicate the onset of the cue. In both trial types, the firing rate scaled with the expected reward size (from left to right). Thus, this neuron showed a consistent encoding of information about reward across different trial types, whether the information was provided by visual cues (top) or whether it relies upon memory from the previous trial (bottom)*.

**Figure 5:**
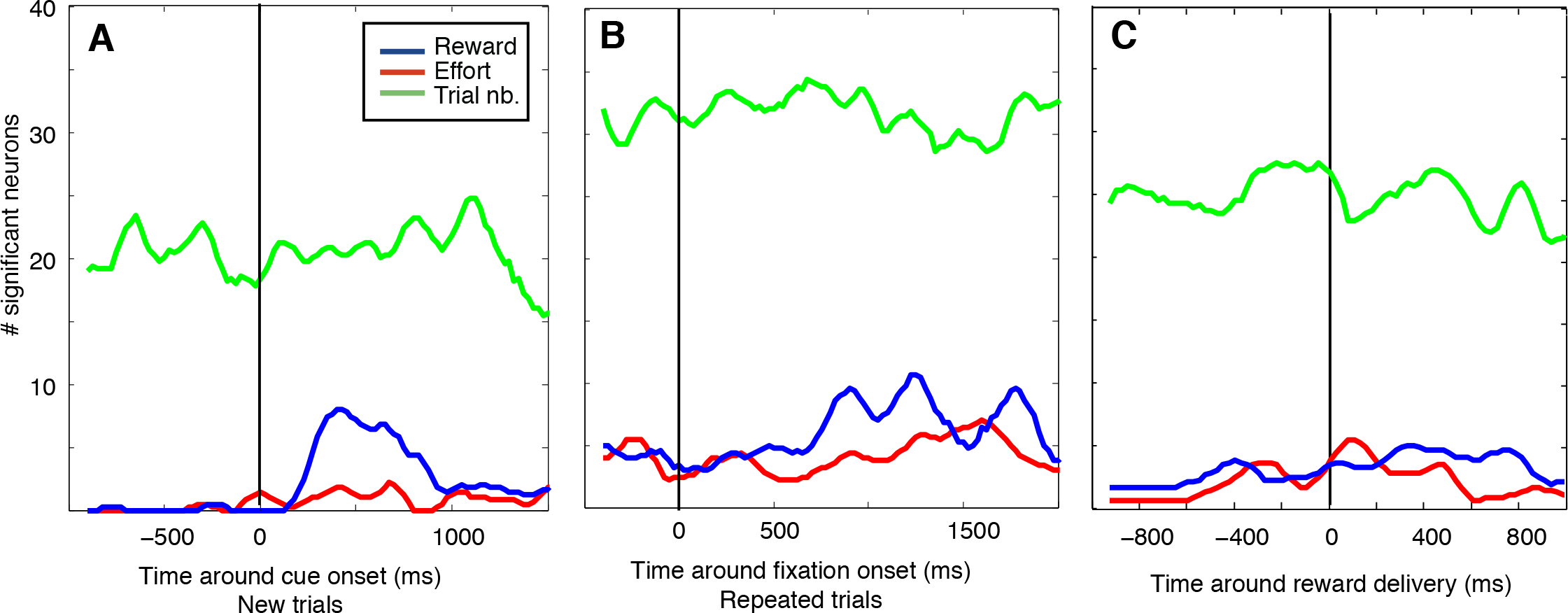
*Modulation of firing by task factors*. *We used a sliding window analysis to measure the sensitivity of all 113 VMPFC neurons to 3 factors: Reward Size (Blue), Effort Level (red) and Trial Number (green). We counted spikes in windows of 200 ms slid by 25 ms around 3 events: the onset of the cue in New trials (**A**), the onset of the fixation point in repeated trials (**B**) and the reward delivery (**C**). For each window, we ran a GLM to estimate the influence of the 3 factors on the firing rate of each unit. We then calculated the number of neuron for which the factor had a significant influence on the firing rate. For each neuron, the effect was considered significant if, in the GLM, the parameter for this factor was significantly different from zero, p<0.05, using FDR correction for multiple comparisons). **A**) Before the onset of the cue in New trials, VMPFC neurons were only encoding progression through the session (trial number). After cue onset, a few neurons started to encode Effort Level, and a few more encoded the expected Reward Size. Note the number of neurons encoding reward and effort is small at least in part because of the correction for multiple comparison, but it is significant because there is a significant increase in the proportion of neurons encoding effort and reward compared to the pre-cue period, when the information was not yet available. The proportion of neurons encoding Trial Number remained high throughout this period. **B**) Around the onset of the fixation point in repeated trials, monkeys could predict the upcoming Reward Size and Effort level before cue onset, based on memory and knowledge about the task structure. A few VMPFC neurons encoded that information. The small increase in the proportion of neurons encoding reward size more than 600 ms after the fixation point is related to the onset of the visual cue, which provide direct information on the trial. Again, a relatively high proportion of neurons encode Trial Number, a proxy for fatigue and satiety that accumulate during the course of the session. **C**) Around reward delivery (about 500 ms after action onset), there was a constant encoding of the 3 factors by VMPFC neurons, with still a greater sensitivity to Trial Number compared to Reward Size and Effort level*.

First, for these 3 task events, a greater proportion of VMPFC neurons encoded the factor Trial Number, which captures the slow effects of fatigue and satiety, than to the information about Effort and Reward. Second, a small but significant proportion of VMPFC neurons encoded the amount of expected reward at the cue onset in New trials (Figure 5A). In the same conditions, we hardly found any VMPFC neuron that was sensitive to the visual information about upcoming effort. In Repeated trials, an equally small but significant proportion of neurons encoded these 2 factors (Figure 5B). An equivalent proportion of VMPFC neurons also encoded these factors around the trial outcome (Figure 5C). In short, VMPFC neurons were more sensitive to advancement through the session (Trial Number) than to information about Reward and Effort. The encoding of these 2 factors engaged a small but significant portion of the population, and VMPFC neurons seemed more sensitive to information about Reward than Effort, especially when it was provided by visual cues.

As shown on figures 3 and 4, the encoding of reward size by VMPFC neurons was relatively consistent over task epochs. We examined more systematically the consistency of encoding of each of these 3 factors (Effort, Reward and Trial Number) across task epochs (Figure 6). Practically, we measured the correlation between estimated regression coefficients of all 113 neurons across 3 epochs of the task: onset of the fixation point, onset of the cue and trial outcome. Overall, Reward Size and Trial Number were encoded reliably across task epochs, but Effort Level was not. More specifically, as shown on figure 3 and 6A (middle), the regression coefficients for Reward at cue onset were significantly correlated with the Reward coefficients estimated at trial outcome (reward delivery). Note that the correlation between Reward coefficients at the cue and the action onset was also significant (p<0.05, data not shown). Thus, the encoding of reward levels by individual VMPFC neurons was very consistent across the different epochs of a trial. In addition, Reward Size was reliably encoded across trial types: the coefficients for Reward were significantly correlated between the onset of the fixation point in repeated trials and the onset of the cue in New trials (Figure 4 for an example neuron; population analysis on Figure 6, center). Using the same approach, we found a similar consistency in the encoding of Trial Number across the different task epochs and across the different trials, in line with the idea that this other factor is encoded reliably in the firing of individual VMPFC neurons (Figure 6, right). We used the same approach to measure the correlation between regression coefficients for the factor Effort across different task epochs but in that case there was no significant correlation (Figure 6, left; all p values > 0.05). This is coherent with the limited sensitivity of VMPFC neurons to information about physical effort compared to Reward and Trial Number.

**Figure 6:**
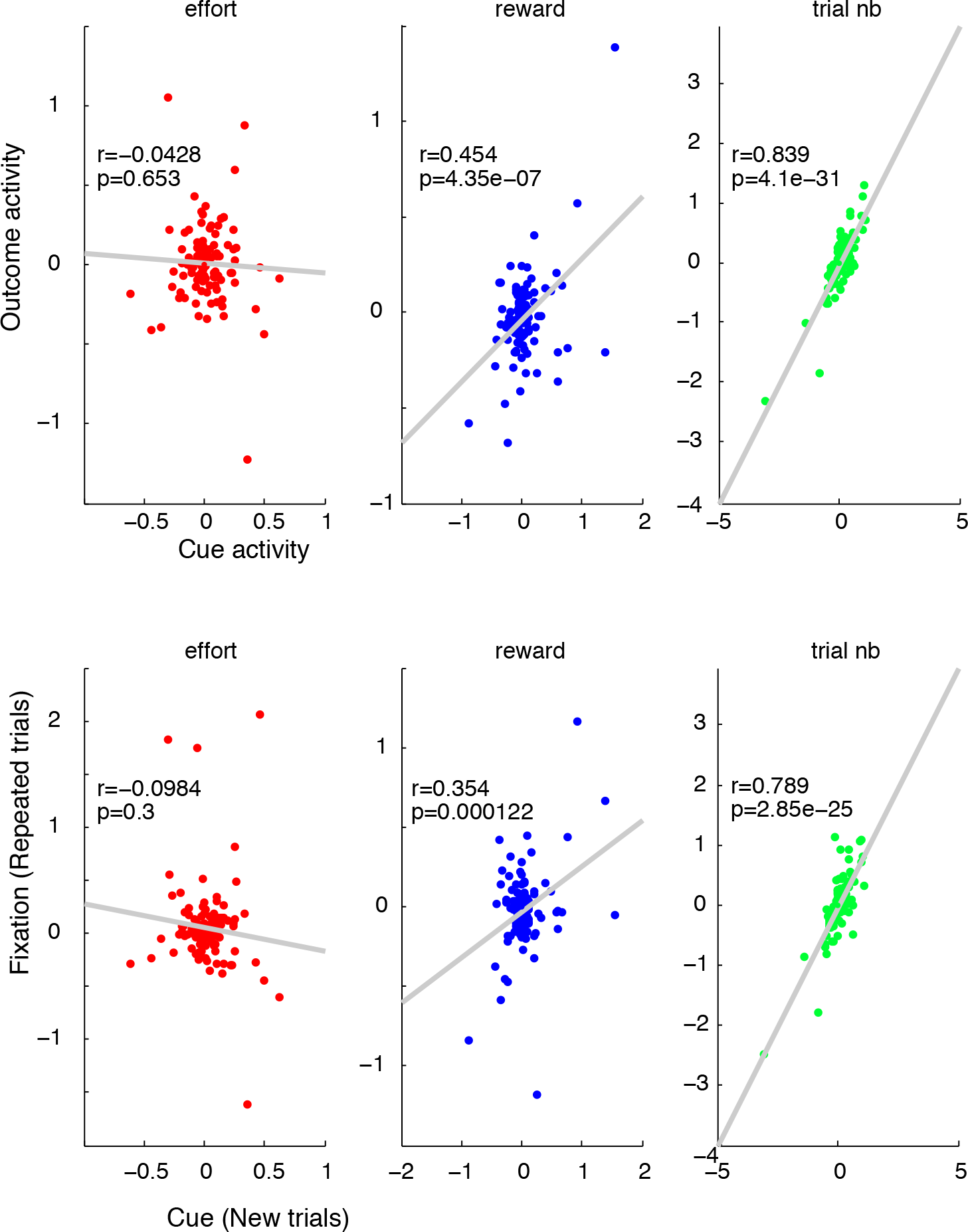
*Consistency of encoding across epochs within and between trials*. *For each neuron, counted spikes in several epochs of the task: At the cue onset and at the outcome in correct trials (**Top**); at the onset of the fixation point in repeated trials and at the onset of the cue in New trials (**bottom**). For each epoch, we ran a GLM analysis to explain firing rate with a constant term plus 3 factors: Effort Level, Reward Size and Trial Number. We estimated the parameters for each of these factors in several epochs of the task and examined the relationship between parameter estimates for the same factor across several epochs of the same trial types (**top**) and between trial types(**bottom**). **Top**: correlation between parameter estimates for effort (left), reward (middle) and trial number (right). For each panel, the x axis corresponds to the activity at the cue and the y axis corresponds to the activity at the outcome, measured in New correct trials. The significant positive correlation between parameter estimates for Reward and Trial Number between these 2 epochs indicates that the encoding of these paramteters by VMPFC neurons is consistent between the onset of the cue and the outcome delivery. **Bottom**: correlation between parameter estimates for effort (left), reward (middle) and trial number (right). For each panel, the x axis corresponds to the activity at the cue in New trials and the y axis corresponds to the activity at fixation point in Repeated trials. The significant positive correlation between parameter estimates for Reward and Trial Number indicates that the encoding of these parameters by VMPFC neurons is consistent between these two epochs in two types of trials*.

To assess the sensitivity of individual VMPFC neurons to information about multiple task factors, we measured the correlation between estimated regression coefficients for pairs of factors across all 113 neurons. None of these correlations were significant (all p values> 0.05). Altogether, these results suggest that Reward level and Trial number induce a coherent firing pattern for individual neurons across trial types and task events, yet their influence on firing is heterogenous and independent across the population of VMPFC neurons.

#### VMPFC activity and willingness to perform the task

After measuring the relation between VMPFC activity and task factors, we examined its relation with behavior. We focused on the willingness to perform the task on a trial by trial basis by measuring the relation between neural activity and choices to perform the action, or not.

We first examined the relation between firing and choices in discrete windows around specific task events, in different subset of trials: cue onset in New trials and fixation point onset in Repeated trials. At the onset of the cue in New trials, only 2/113 neurons encoded choices. However, as shown on figure 1, choices in these conditions displayed little variability and virtually no influence of Trial Number. As discussed above (section 'Behavior'), monkeys have a strong bias for performing the action in New trials and whatever its source, this strong positive bias probably accounts for the lack of modulation by task factors, for both choices and neuronal activity. The proportion of neurons encoding choices was greater at the onset of the fixation point in Repeated trials (n=28/109), where choices were both more variable and more sensitive to Reward, Effort and Trial Number. There are many features that differ between New and Repeated trials, including the presence of the cue and the history of recent events, so drawing a direct comparison between neural activity in these conditions is difficult. But it is safe to conclude that a significant proportion of VMPFC neurons encoded the willingness to engage in the task and perform action to get the reward when monkeys displayed enough variability in their choices.

Altogether, this indicates that VMPFC neurons provide a temporally stable representation of information about reward and trial number. When monkeys readily use this information to adjust their willingness to perform the task (in Repeated trials), VMPFC also reflects the resulting course of action. This indicates that VMPFC activity does not reflect simple sensory-motor processes. Rather, VMPFC neurons were mobilized when monkeys adjusted their behavior based on contextual factors. Next, we decided to explore the slow dynamics of the relation between VMPFC firing and the animals’ willingness to work.

#### Slow modulation of VMPFC activity and willingness to work: influence of previous trials.

As shown for a representative example on Figure 7, the dynamics of the relation between VMPFC and the willingness to engage in the task was relatively slow, developing over seconds and encompassing several trials. Since the previous analysis was based on subset of trials, it could not capture the slow dynamics of that relation. To quantify the slow and continuous modulations of the willingness to work and of VMPFC activity over successive trials, we measured the influence of previous trials on two variables: the choice to perform the task or not and the firing at the onset of the fixation point. Critically, we included every trial in the analysis and we evaluated the influence of previous trials over and above current levels of Reward, Effort and Trial Number.

**Figure 7:**
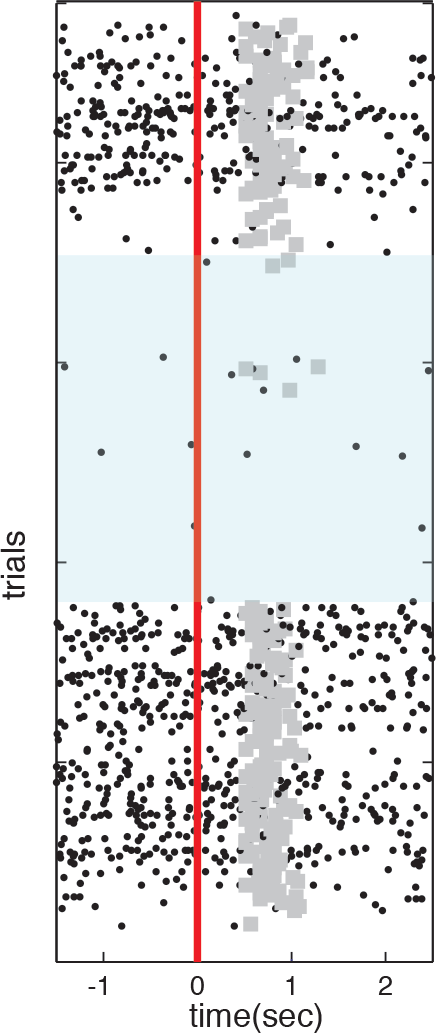
*Slow encoding of the engagement in the task in the VMPFC*. *Activity of a single VMPFC neuron around onset of the fixation point (black vertical line) with trials sorted in chronological order (first at the top) Grey dots represent the onset of the cue, in trials were the monkey did not break fixation before it appeared, i.e. when it chose to engage in the task at least until cue onset. The firing of the neurons was clearly modulated by the willingness of the monkey to perform the task: At the beginning of the session, the monkey was willing to work and the neuron was firing robustly. Then the activity of the neuron decreased and a few trials later the monkey completely stopped working for a relatively long period (shaded area). Note that the few trials initiated in the middle of this long break were associated with an increase in firing. Later on, the animal resumed working at the same time as the neuron increased its firing rate. At the very end of the session (at the bottom of the plot), the firing of the neuron decreased again a few trials before the animal stopped working*.

For the behavior, we used a simple logistic regression to account for the choice to perform the task in any trial n with a weighted linear combination of the following regressors: Reward, Effort, and Trial Number at trial n, as well as choices at trial n-x, with x ranging from 1 to 35. This allowed us to measure the relative weight of previous choices as a function of the distance × (in trials) between previous and current choice. The willingness to perform the task in a given trial was related to the behavior up to 20 trials in the past, over and above the positive influence of current Reward and the negative influence of Effort levels and Trial Number (Figure 8A).

**Figure 8:**
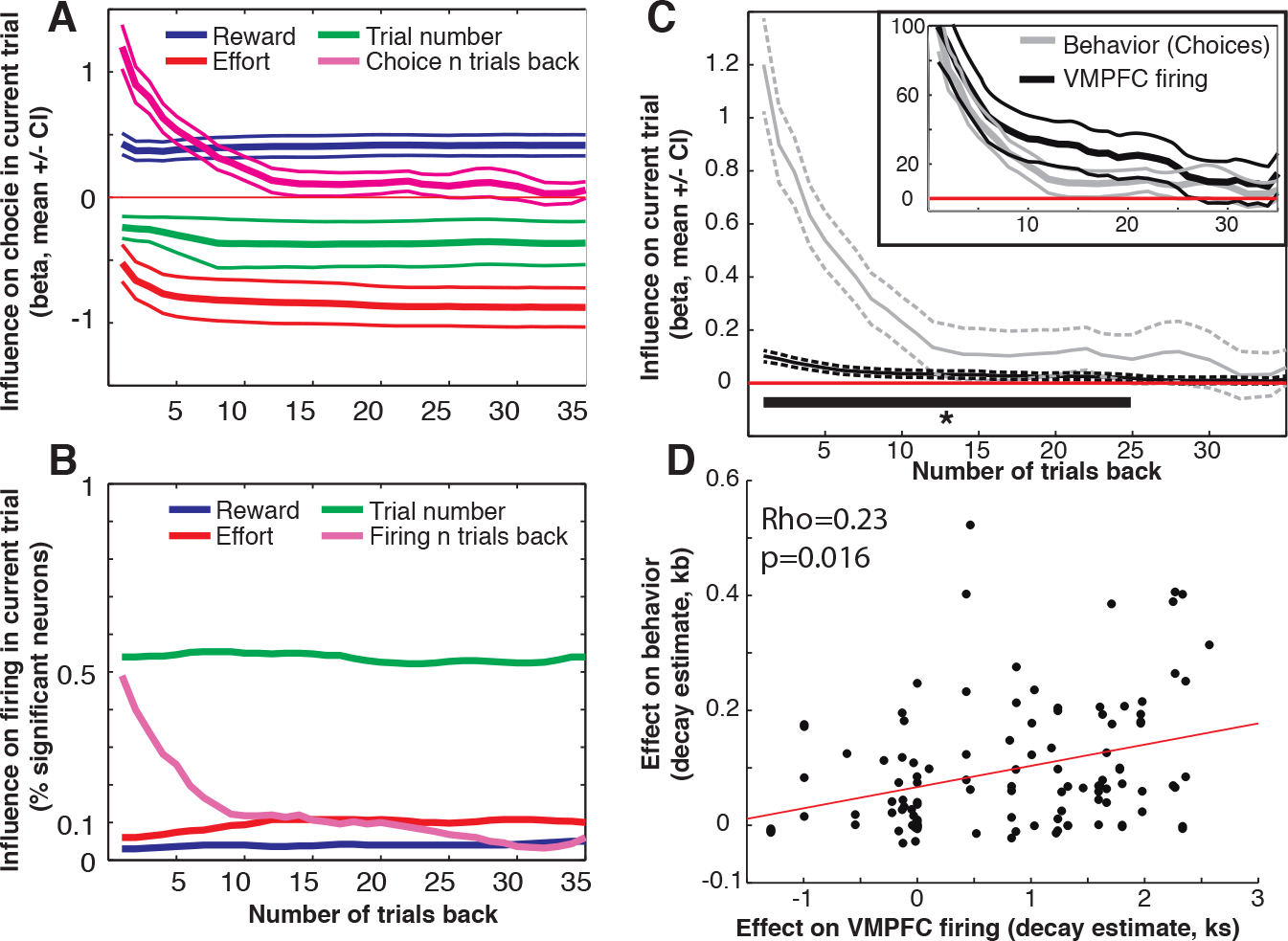
*Influence of past trials on willingness to work and VMPFC firing.* ***A**) Relative influence of task parameters and previous choices on engagement in the task. We used a logistic regression to estimate the modulation of the willingness to perform the task as a function of task parameters (Reward, Effort and Trial Number) in the current trial as well as choices in past trials, as a function of the number of intervening trials between current and past choice (x axis). We included all trials in the analysis. The lines represent the mean and confidence interval of the parameters estimates (Betas) describing the relative influence of Trial Number (green), current Reward (blue) and Effort (red) levels and choice x trials back. In line with earlier analysis, monkeys did take into account information about Reward, Effort and Trial Number to adjust their willingness to engage in the trial at the onset of the fixation point. There was a strong positive influence of choice in recent trials (x<5), indicating a tendency to repeat previous choices, over and above all the other parameters. This influences decreases as the number of intervening trials between current and past choice increases. **B**) Relative influence of task parameters and activity in recent trials on VMPFC firing. We used a GLM to estimate the relative influence of firing in previous trials and task parameters (Reward, Effort and Trial Number) on firing in the current trial (y axis), as a function of the number of intervening trials between current and past trial (x axis). We included all trials of all 113 neurons in the analysis, and examined the activity at the onset the fixation point. The lines represent the fraction of neurons showing a significant effect of the corresponding parameter, after correction for multiple comparisons. In line with previous analysis, there were more neurons showing a significant effect of Trial Number compared to Reward and Effort. On top of these effects, VMPFC neurons displayed a strong sensitivity to activity in previous trials (pink line). In other words, activity at the fixation point is strongly correlated over many successive trials, in line with the idea that the firing of VMPFC changes slowly during sessions, over and above task parameters. **C**) Dynamics of the influence of past trials on behavior and VMPFC firing. For both choices and VMPFC activity, in each session, we extracted the beta values of the regression described in A and B, respectively, and for each of the distances between current and past trial. The lines correspond to the mean and confidence interval (2 standard errors) for each of the distances, for both the willingness to perform the task (grey) and VMPFC firing (black). The data was smoothed for representation purposes. The inset shows the same distributions after values were normalized to 100%, to facilitate the comparison. A second level analysis on these distributions (t-test, with correction for multiple comparison using False Discovery rate procedure) revealed that, for both choices and spike counts, the influence of previous trials was significant for up to 25 trials between past and current trials. **D**) Influence of past responses: relation between behavioral and neurophysiological effects. We evaluated the shape of the relation between past and current trials as a function of the number of intervening trials by fitting the data in each session with a exponential decay function (see text for further details). We estimated the parameter k by fitting the relation between response in trial n as a function of x trials back with the function: response=a.exp (-k.x). For each recording session, we repeated the procedure for the behavioral response (to obtain a variable kb) and for VMPFC spiking activity (to obtain a variable ks). This plot shows the relation between kb and ks variables obtained in all 113 recording sessions. The significant correlation between the two indicates that the dynamics of the impact of previous trials on behavior and VMPFC activity are correlated across the different recording sessions*.

To estimate the influence of firing in previous trials on VMPFC activity, we used a regular GLM to account for spike count at the onset of the fixation point at trial n based on task factors at trial n as well as spike count at trials n-x. We also observed a significant impact of activity in past trials on the firing of VMPFC neurons at trial onset (Figure 8B). Note that we quantified the effect at the population level by counting the number of significant neurons because since there was no tendency for VMPFC neurons to encode task parameters in a systematic positive or negative fashion, the average beta values for Reward, Effort and Trial Number was not significantly different from zero (second level analysis with t-tests, all p values >0.05). By contrast, as shown on Figure 8C (black line), the relation between firing in current versus past trials was clearly positive, indicating that the firing of VMPFC neurons was relatively consistent over several successive trials, over and above changes in firing induced by changes in Reward, Effort or Trial Number.

On average, the influence of previous trials on behavior and VMPFC firing showed a very similar temporal profile (Figure 8C), with a strong influence of responses occurring up to 20 trials before on behavioral and neuronal responses in a given trial. Given that similarity, we examined the possibility that this slow changes in behavior and VMPFC activity were directly related, on a session by session basis. For both behavior and VMPFC activity, we measured the influence of past responses as a function of the distance between past and current response (ranging from 1 to 35) in each recording session. Note that the average curve for all sessions is available on panel C, for both behavior and neuronal responses. For each measure (behavioral and neuronal responses) and each session, we fitted the influence of previous responses with a simple exponential function using a variational Bayes method, using the VBA toolbox in Matlab (Daunizeau et al, 2014). Practically, we calculated the optimal parameters a and k describing the function y=a.exp(-k*x), where y is the effect size of the influence of past responses, over and above task parameters (measured using the beta coefficient in the GLM described before) and x is the distance between past and current response, ranging from 1 to 35 trials. After extracting this couple of parameter a and k for both spiking (a_s_ and k_s_) and behavior (a_b_ and k_b_) in each session, we measured the correlation between parameters describing behavioral and neuronal responses across all 113 recordings. There was a significant positive correlation between both pairs of parameters (a_s_ and a_b_: rho=0.33, p=3.410^−4^; k_s_ and k_b_: rho= 0.23; p=0.016; Figure 8D). Thus, across recording sessions, there was a significant relation between the influences of past trials on the monkeys’ willingness to perform the task and VMPFC firing.

#### Slow modulation of VMPFC activity and willingness to work: beyond task events.

Finally, we reasoned that if the firing of VMPFC neurons was encoding the willingness to engage in the task in a continuous fashion, it should reflect fluctuations in work rate (number of trials performed in a given duration) at time scales longer than individual trial events and even longer than the duration of a trial (about 3-4 seconds). To explore more systematically the relation between VMPFC activity and engagement in the task across time scales, we compared the relation between firing rates and work rate by splitting each session into successive time windows ranging from 2 to 100 seconds (Figure 9). This analysis follows on the idea that the encoding of task factors was stable across task events and trial types, allowing for investigation of average activity over long recording periods. Note that the statistical power decreases when the time window is prolonged, as there are fewer data points to assess the correlation. Consistently the strength of the correlation between firing rate and Work Rate (grey line) displayed a monotonic decrease over window sizes. Yet a large number of neurons significantly encoded Work Rate even at longer time scales.

**Figure 9:**
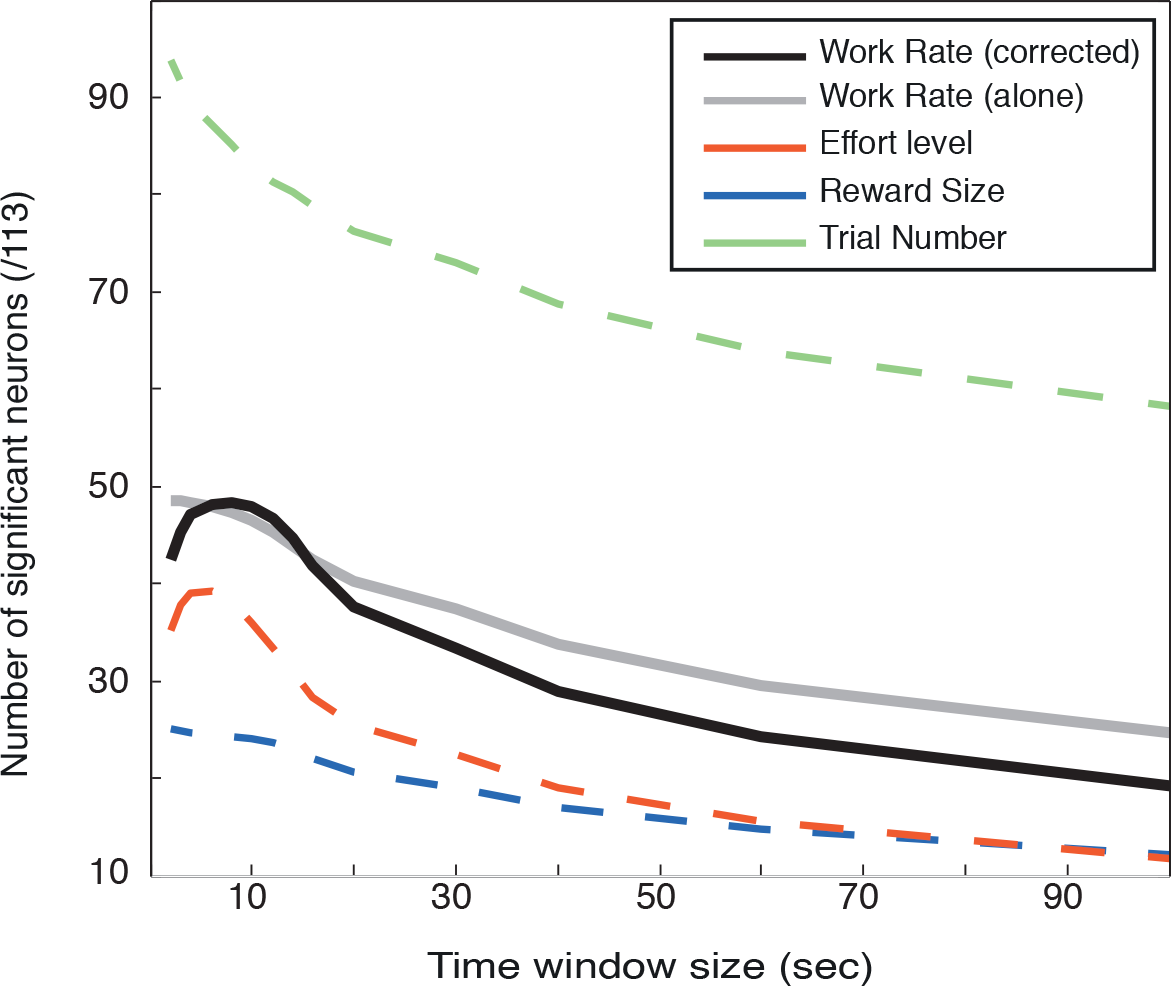
Slow encoding of the engagement in the task in the VMPFC. *To examine the encoding of Action Value at different time scales, we split each recording sessions into successive time windows of variable sizes (from 2 to 100 seconds, x axis). For each window size, we measured the relation between the number of trials performed (Work Rate) and firing rate of VMPFC neurons. The Y axis indicates the number of neurons displaying a significant relation with Work Rate. We used a GLM to predict neuronal firing of each neuron across all the windows of the session using either Work Rate alone (grey) or Work Rate corrected for the effect of the task factors (black). The broken lines indicate the number of neurons displaying a significant modulation of firing by these 3 factors (Reward size, blue, Effort level, red, and Trial Number, green), which we used for correcting firing rate and Work Rate across all window sizes. There is a significant relation between VMPFC activity and willingness to perform the task in a large fraction of neurons, over and above the task parameters, and it can be captured even outside of tasks events, and at slower time scale compared to the pace of the task*.

To verify that this correlation was not simply due to the influence of Reward, Effort and Trial Number, which all vary at the scale of trial duration (3-4 s), we conducted a second analysis where we corrected Work Rate for the effects of these factors by including them as co-regressors in the model to predict firing rate. Practically, we used a GLM to evaluate the impact of 4 variables on VMPFC activity: work rate (corrected), Reward, Effort and Window Number (see methods). The sensitivity of VMPFC neurons to the work rate increased for windows of 2 to 10 seconds and decreased for wider windows, but the number of neurons showing a significant effect remained above chance level (n=16/113). Note that the proportion of neurons showing a positive and negative relation between firing rate and work rate were equivalent, so that overall, there was no global change in the firing rate of the population in relation to the slow changes in work rate. In short, for a large fraction of VMPFC neurons, the relation between spiking activity and willingness to work was strong and reliable enough over time to appear when analyzed at a time scale much longer than task events.

## Discussion

In summary, VMPFC neurons coherently encoded information about upcoming reward and progression through the session across distinct task epochs. The encoding of the upcoming effort was weaker and less coherent, but it also affected VMPFC firing at a slower time scale. The activity of VMPFC neurons was strongly related to the willingness to engage in the task, and it captured the influence of multiple factors on behavior, beyond individual task events. Indeed, VMPFC neurons seem to monitor information over time to encode the value of the current course of action in a continuous fashion, over and above the specific task parameters. Thus, VMPFC neurons can continuously integrates internal and external information about potential costs and benefits, to determine the global willingness to engage in a goal-directed behavior.

### 1) <u>Encoding of Reward and Effort by VMPFC neurons</u>

Even if the proportion of neurons encoding Reward Size was relatively limited compared to Trial Number, it was not negligible since it survived corrections procedures. Moreover, the encoding of the expected reward size was consistent across task epochs, whether reward was experienced directly (at trial outcome), announced by visual stimuli (at cue onset) or just predicted by memorized information (at fixation point). This reinforces and extends recent neurophysiological studies in monkeys showing that VMPFC neurons reliably encode reward information (Monosov and Hikosaka, 2012; Strait et al., 2014).

Our task includes an effort component, which is critical for understanding how the VMPFC contributes to balancing energetic costs and benefits for decision making. But as seen in recent human neuroimaging data (Croxson et al., 2009; Prevost et al., 2010; Skvortsova et al., 2014), VMPFC activity was less sensitive to information about effort, especially when it was provided by visual stimuli. This is in line with the idea that effort processing engages more strongly the anterior cingulate cortex, as was initially found in rats and more recently in monkeys (Walton et al, 2002, Rudebeck et al, 2006, Hosokawa et al., 2013). Interestingly, we and others also observed a higher sensitivity to reward versus effort in the activity of dopaminergic neurons, which, like the VMPFC, are thought to be important for reward processing (Gan et al., 2010; Pasquereau and Turner, 2013; Varazzani et al., 2015).

Note, however, that in our experiment a significant number of VMPFC neurons did encode effort when information was provided by contextual information, both on a trial by trial basis (Figures 5B and 8B) and when we analyzed activity at slower time scale (see Figure 9). Thus, the difference between reward and effort encoding was most visible following cue onset in new trials, which corresponds to the situation examined in human fMRI experiments (Croxson et al., 2009; Prevost et al., 2010; Skvortsova et al., 2014). It is possible that this phasic activation to expected reward represents a reward prediction error, since before new trials the expectation is always the same, such that reward level and reward prediction error are confounded at this epoch. In contrast, tonic encoding of reward and effort levels engaged an equivalent proportion of VMPFC neurons. This was the case both when the information was already known (in Repeated trials) or when taking larger time windows that provide more robust estimates. It remains possible that the continuous coding of effort level represents a subjective effort estimate, which would be more relevant to decision making than the objective amount of force to exert, which is more important for motor control. This subjective estimate might relate to reward probability, if the effect of effort level on the choice not to perform the task is taken into account. Alternatively, subjective effort might correspond to the discomfort induced by action execution, which would be integrated as a (negative) sensory feedback in brain regions representing the outcome space and not the action space. This idea that information about effort has a different meaning across task conditions could also account for the lack of coherence in the encoding of this parameter across task epochs, compared to reward size and trial number (Figure 6). But beyond these limitations, VMPFC neurons are more sensitive to effort when its influence can be integrated over time, which is in line with the general idea that VMPFC neurons integrate information relevant for guiding behavior over a relatively slow time scale.

There was no correlation among the sensitivities of individual VMPFC neurons to the three task factors (Reward, Effort and Trial Number). This is in contradiction with recent studies reporting a significant correlation between the estimated parameters capturing the influence of distinct task factors (Abitbol et al., 2015; Strait et al., 2014). But this is coherent with studies indicating that the ability to 'multiplex' information is significantly stronger in more dorsal and posterior regions of the medial prefrontal cortex (Hosokawa et al., 2013; Kennerley and Wallis, 2009). Thus, multiplexing is probably not the most critical feature of VMPFC neurons. Note that the absence of correlation does not imply that the different task factors were represented in different populations of neurons. It simply means that the code might be more complex than initially thought, as the weights of the different factors were independently distributed over neurons.

### 2) <u>Sensory vs contextual information</u>

Another key feature of VMPFC neurons is their strong sensitivity to contextual information, compared to information provided by sensory stimuli.

First, in line with our previous work, a very large fraction of VMPFC neurons encoded the progression through the session (Trial Number), which is considered a proxy for fatigue and/or satiety, was also very reliable across task events and trial types and it is in line with our previous work (Bouret & Richmond, 2010). Note that the much stronger sensitivity of VMPFC neurons to Trial Number compared to Reward and Effort level was at odds with the relatively balanced influence of these factors on behavior (Figures 1C and 5), which confirmed the idea that VMPFC neurons were particularly sensitive to contextual information (here in the physiological domain).

Second, even if it was relatively small, the percentage of VMPFC neurons encoding Reward and Effort levels was as high or higher in conditions when the information relied on memory (Repeated trials) compared to when it relied upon visual stimuli (New trials). It is difficult to identify the nature of the memorized information in this task, and it presumably includes recent informations about task conditions, recent actions, physiological state and knowledge about the task. But irrespectively of the nature of that information, it is not provided by direct sensory cues and it has a strong influence both on behavior and VMPFC firing. The sensitivity of VMPFC neurons to that contextual information is compatible with lesion studies in rats and monkeys emphasizing the specific role of medial orbitofrontal cortices in computing action value using both observable and unobservable information (Noonan et al, 2010; Bradfield et al., 2015).

This feature, together with the coherent coding over task events, seems much more prominent in the VMPFC compared to more lateral regions of the ventral prefrontal cortex (Abitbol et al., 2015; Bouret and Richmond, 2010). In the OFC the encoding of information related to reward appears more specific and better locked in time to critical events (Blanchard et al., 2015; Howard et al., 2015; Wilson et al., 2014). This is also in line with the imaging literature in humans, which emphasizes the critical role of the VMPFC, but not the OFC, for the representation of subjective value (Bartra et al., 2013; Boorman et al., 2013; Chib et al., 2009; Clithero and Rangel, 2014; Lebreton et al., 2009). The subjective value represented in the VMPFC integrates the value of information bits that are encoded in distinct brain regions, notably when they are not directly observable (Barron et al., 2013; Benoit et al., 2014; Hare et al., 2010). Finally, this is coherent with our past work describing the complementary roles of OFC and VMPFC in stimulus-bound vs context-dependent reward processing, respectively (Bouret and Richmond, 2010).

### 3) <u>Slow relation between VMPFC activity and willingness to work</u>

Another important feature of VMPFC activity in this task is the relatively slow time scale with which it integrated information about the task or about the behavior. This is in line with our recent work showing a strong relation between pre-stimulus activity and the subjective value of visual cues, both using single units in monkeys and fMRI in humans (Abitbol et al., 2015). Here, as discussed above, we showed that the encoding of Reward Size and Trial Number was very coherent over time. We also observed a strong autocorrelation in the firing of VMPFC neurons across successive trials, over and above the influence of reward, Effort and Trial Number. In addition, the autocorrelation in the activity of VMPFC neurons mirrored the strong autocorrelation in behavior (Figure 8). At the behavioral level, this tendency to maintain the same behavior over up to 20 successive trials, over and above task parameters, suggests that animal go through series of states. These behavioral states were characterized by distinct levels of engagement in the task. The slow dynamics of VMPFC neurons was directly related to these slow fluctuations in the animals’ willingness to work (Figures 7-9). Thus, VMPFC neurons might be directly involved in the encoding of these motivational states, defined by a global willingness to perform reward directed actions over and above task relevant information. This feature might be important for understanding the implication of the VMPFC in mood and its disorder such as depression (Ressler & Mayberg 2007; Rutledge et al, 2014).

This ability to encode the willingness to work over relatively long time scales, rather than discrete sensory-motor operations, is consistent with the notion that value coding in the VMPFC is automatic. Indeed, the encoding of value in the VMPFC appears even when subjects are not engaged in a valuation task such as choice or rating (Harvey et al., 2010; Lebreton et al., 2009; 2015; Levy et al., 2011). Here the effort task was imperative and did not include any explicit choice phase with different responses corresponding to different options, and yet VMPFC neurons continuously encoded the willingness to work. The fact the VMPFC automatically and continuously encodes the willingness to perform the task might also explain why this region has been implicated in the so-called default mode network (Raichle and Snyder, 2007). Indeed this network is typically observed in contrasts between blocks of effortful cognitive tasks and resting periods during which subjects may mind-wander around the topics they like.

### 4) <u>Conclusion & perspectives</u>

Altogether, this work is directly related to the emerging idea that the VMPFC computes outcome value based on contextual/memory information through a direct interaction with hippocampus and associated cortices (Aminoff et al., 2013; Barron et al., 2013; Benoit et al., 2014; Clark et al., 2013; Lebreton et al., 2013; Lin et al., 2015; Noonan et al., 2010; Peters and Büchel, 2010, Brown et al, 2016). This ability to adjust behavior as a function of memorized information is probably critical for primates, which generally forage for fruits in complex and variable environments (Noser and Byrne, 2015; Cunningham and Janson, 2007; Janmaat et al., 2013; 2011). Thus, the development of this system involving the VMPFC in primates might be directly related to a strong ecological pressure to integrate information about costs and benefits over time and compute the willingness to engage in reward-directed actions based on both immediate and memorized information. One of the challenges ahead is to understand how distinct physiological and ecological constraints have shaped the relative development of these functions and the associated structures across primate species.

## Acknowledgments

The authors declare no competing financial interests. We would like to thank Jean Dau-nizeau for helpful comments on data analysis. We would like to thank Morgane Monfort, Serban Morosan and the personnel from the ICM primate facility for assistance with surgery and veterinary procedures. This work was supported by a Starting Grant founded by the European Research Council (ERC-BioMotiv). C.V. received a Ph.D. fellowship from the “Ècole doctorale Frontieres du Vivant” (FdV).

